# scMARK an ‘MNIST’ like benchmark to evaluate and optimize models for unifying scRNA data

**DOI:** 10.1101/2021.12.08.471773

**Authors:** Swechha, Dylan Mendonca, Octavian Focsa, J. Javier Díaz-Mejía, Samuel Cooper

**Affiliations:** Phenomic AI

## Abstract

Today’s single-cell RNA analysis tools provide enormous value in enabling researchers to make sense of large single-cell RNA (scRNA) studies, yet their ability to integrate different studies at scale remains untested. Here we present a novel benchmark dataset (scMARK), that consists of 100,000 cells over 10 studies and can test how well models unify data from different scRNA studies. We also introduce a two-step framework that uses supervised models, to evaluate how well unsupervised models integrate scRNA data from the 10 studies. Using this framework, we show that the Variational Autoencoder, scVI, represents the only tool tested that can integrate scRNA studies at scale. Overall, this work paves the way to creating large scRNA atlases and ‘off-the-shelf’ analysis tools.

## 1 Background

Single-cell RNA sequencing (scRNA) studies can dissect tissues and biological processes with exquisite detail. An explosion in scRNA studies has also emerged from rapidly falling costs associated with generating scRNA data [1], Fig. 1. Typically, scRNA analysis focuses on individual datasets. An ever-growing number of published studies creates an opportunity for wider integrated analysis [2]. Yet, the challenge of combining data from multiple disparate scRNA studies remains [3]. Accurate machine-learning (ML) models that can integrate multiple scRNA studies offer the opportunity to generate robust single-cell transcriptional atlases. Yet we lack the high-quality benchmark datasets and evaluation frameworks needed to effectively test how well models align many studies [4, 5, 6, 7, 8].

**Figure 1:**
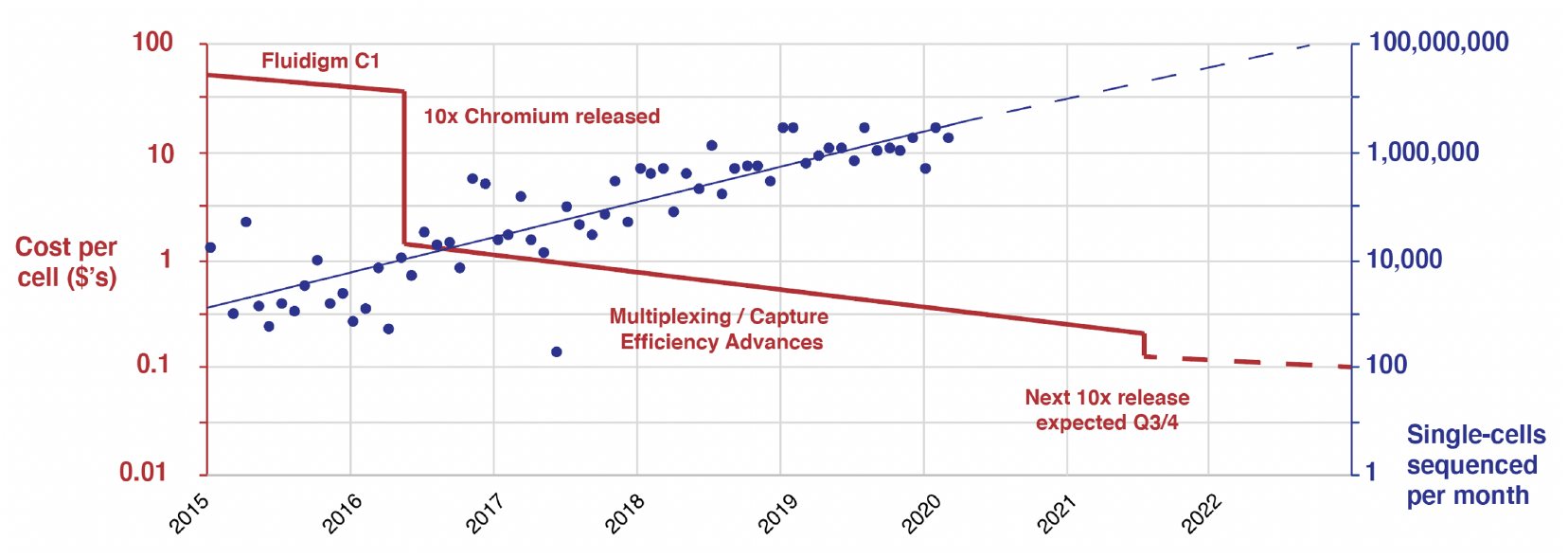
scRNA studies are on the rise: The number of cells published per month is increasing exponentially, as per Svensson et al. [1]. We are also seeing the sequencing cost per cell fall rapidly; internal analysis, key reference points used, e.g., Zhang et al. [13]

Some of the most significant improvements in machine-learning for image analysis came from improving model performance on high-quality ‘benchmark’ datasets. For example, the MNIST dataset contains 1000’s of handwritten digits (0-9) and led to breakthroughs in machine learning in the 1990s and 2000s, e.g., the development of deep convolutional neural networks [9]. We propose that a lack of good scRNA benchmark datasets stems from two challenges:

1. No clear ground-truth exists, instead cell-types are labelled by authors. Comparisons therefore often rely on statistical performance metrics [10, 11] or cell-type labels given by an alternate experimental method, e.g., CITEseq, themselves subject to bias [12].
2. In scRNA, we want to test the ability of an unsupervised method to reduce a high-dimensional gene space to a low-dimensional ‘cell-type’ space while aligning studies, as this enables new cell-type discovery. Yet this creates a challenge since ground-truth labels are typically used to assess supervised methods.

## 2 A framework for evaluating scRNA model performance

We propose a solution to these challenges whereby (1) We assume that, on average, author labels are correct. Thus, we can test how well models predict author labels on data from held-out studies (Fig. 2A); and (2) We assume that cell types can be easily separated in good cell-type spaces, while separation is challenging in poor cell-type spaces. Therefore, we can use a collection of simple supervised models to evaluate how good a cell-type space is by seeing how well they can label cell types on a held-out dataset (Fig. 2B & 2C).

**Figure 2:**
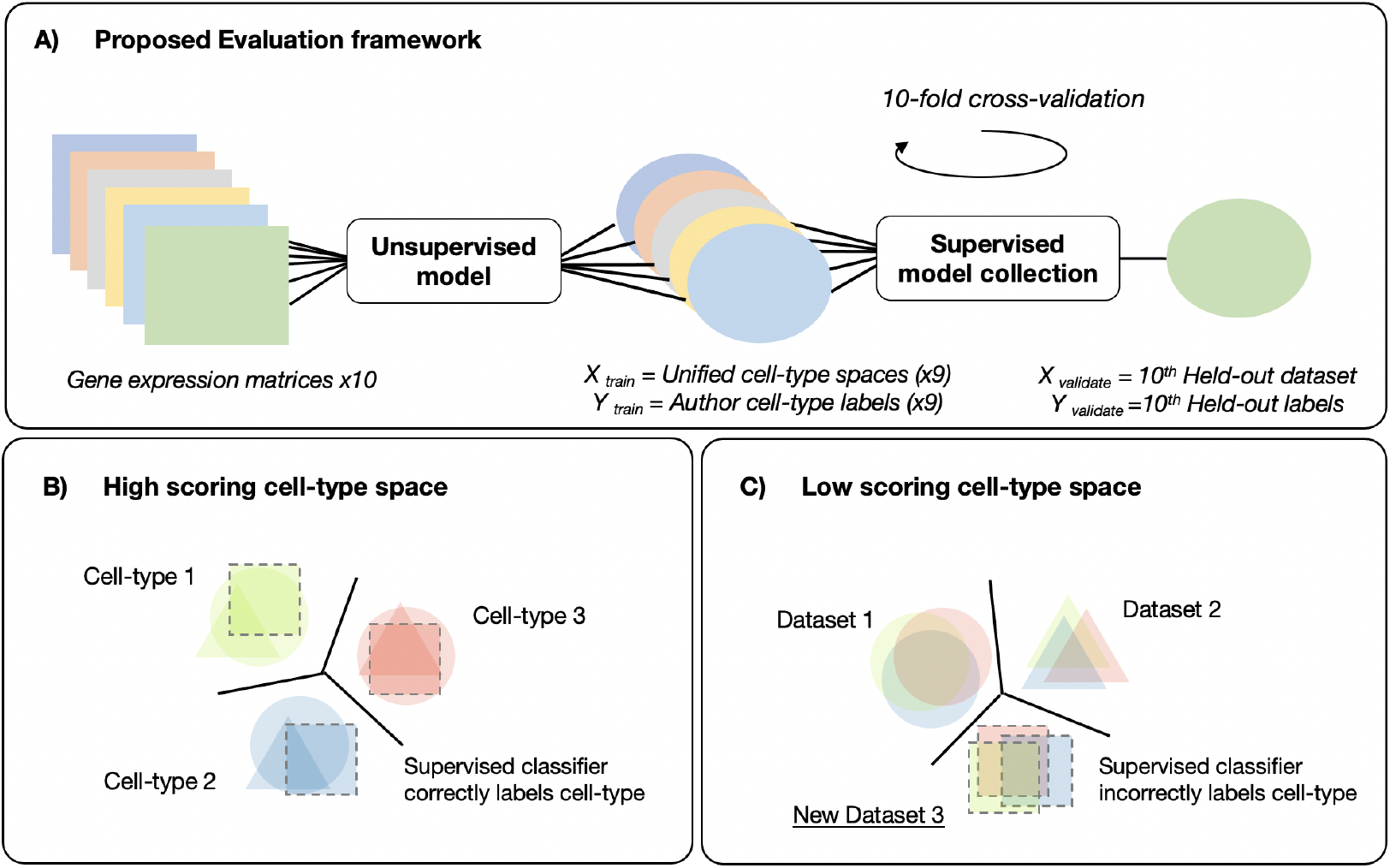
A framework for evaluating model performance at integrating scRNA datasets: A) In our evaluation framework, we use unsupervised models to reduce the complete set of single-cell gene expression matrices into a unified cell-type embedding space. We then train a collection of supervised models to predict author labels from all but one held-out dataset in this unified cell-type space. We then evaluate the performance of the supervised models on the held-out dataset with traditional accuracy metrics, e.g., F1 score; B) We propose that supervised models find it easy to predict cell-type labels in new studies, in high-scoring cell-type spaces, since cell-types are well aligned; C) In low-scoring cell-type spaces, supervised models find it challenging to label cell-types since poor alignment makes it difficult to predict cell-type labels.

Under these assumptions, we can therefore identify top-performing models by (1) Generating a benchmark dataset comprising cells from stratified samples from a set of author labeled scRNA studies, and with harmonized cell-type and gene labels; (2) Using unsupervised scRNA analysis models to reduce gene-space to an aligned cell-type space; and then (3) Evaluating the quality of this cell-type space by training a collection of supervised models to label cell-types on all but a held-out study that we use to assess performance (with cross-validation).

## 3 Results

### 3.1 Generation of the scMARK benchmark dataset

We unified ten scRNA studies (Table S1), taking a random 10,000 cell sample from each study. Our goal was to maintain author cell-type labels as closely as possible, while standardizing labels across studies. Our harmonization process recorded 28 cell-type labels across the studies (Fig. 3A and Table S2). We also harmonized gene naming conventions using the latest versions of the Human Protein Atlas [14], and ENSEMBL [15] databases. Genes were kept that existed at the intersection of at least 2 of the ten studies (Fig. 2B). Of note, Bi et al. 2021, reported vastly more gene identifiers than other studies, as this study also captured non-coding RNA, whil, Azizi et al. 2018 reported less, likely due to the use of the inDrop in this study, vs. the others that used a 10X instrument. Overall, this benchmark dataset represents the first effort to unify studies with author specified ground-truths, as has previously been proposed as a gold standard [8], but until now not addressed owing to the harmonization challenges we solve here.

**Figure 3:**
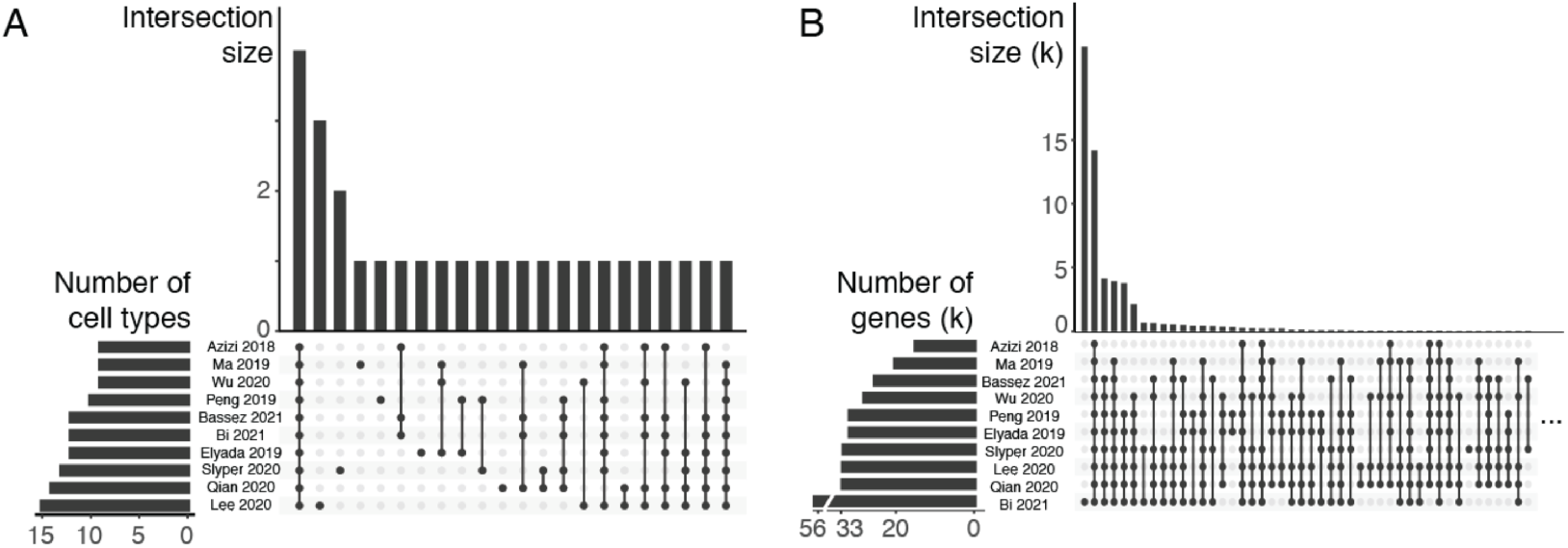
Results of dataset harmonization: A) Only a handful of cell-type labels are annotated across all studies, highlighting variation in author labelings (e.g., T, NK, and B cells). B) At the gene level, a much greater level of overlap between naming conventions was seen, with most genes being present in all studies.

### 3.2 Model performance evaluation

For the unsupervised step of our evaluation framework we tested; (a) Principal Component Analysis (PCA) applied directly to all genes (PCA_*all*_), (b) Reciprocal PCA as per Seurat using the ‘SC-Transform’ data slot (RPCA_*sct*_); (c) Reciprocal PCA as per Seurat using the ‘integrated’ data slot (RPCA_*int*_) [16] (d) PCA applied to a subset of genes selected as described in scmap (PCA_*scmap*_) [17]; and (d) Variational Auto-Encoders (VAEs), implemented as per single-cell Variational Inference models described in Lopez et al. (scVI) [18]. For PCA_*all*_, PCA_*scmap*_ and scVI we used the log normalized gene expression matrices from all 10 studies. Whereas, for RPCA_*sct*_ and RPCA_*int*_ we used the SCTransform normalization. Each unsupervised unified cell-type space was input to four cell-type classifiers: XGboost (XGB), Gaussian Naive Bayes (GNB), K-nearest neighbors (KNN), and Support Vector Machine (SVM) models, which acted as our collection of supervised models to evaluate the cell-type space.

Based on this framework, we found that scVI outperformed other approaches using the scMARK benchmark dataset (Fig. 4A; Table S2). PCA_*all*_ and PCA_*scmap*_ generated results comparable to each other. Finally, RPCA_*sct*_ performed poorly on this benchmark. Importantly, we found minimal variation in the mean F1 score between supervised methods in contrast to the unsupervised integration methods, implying we are successfully evaluating the unsupervised model. Overall, this comparison shows clear differences in performance, validates our two step strategy for testing unsupervised models and shows that scVI stands out above other methods.

**Figure 4:**
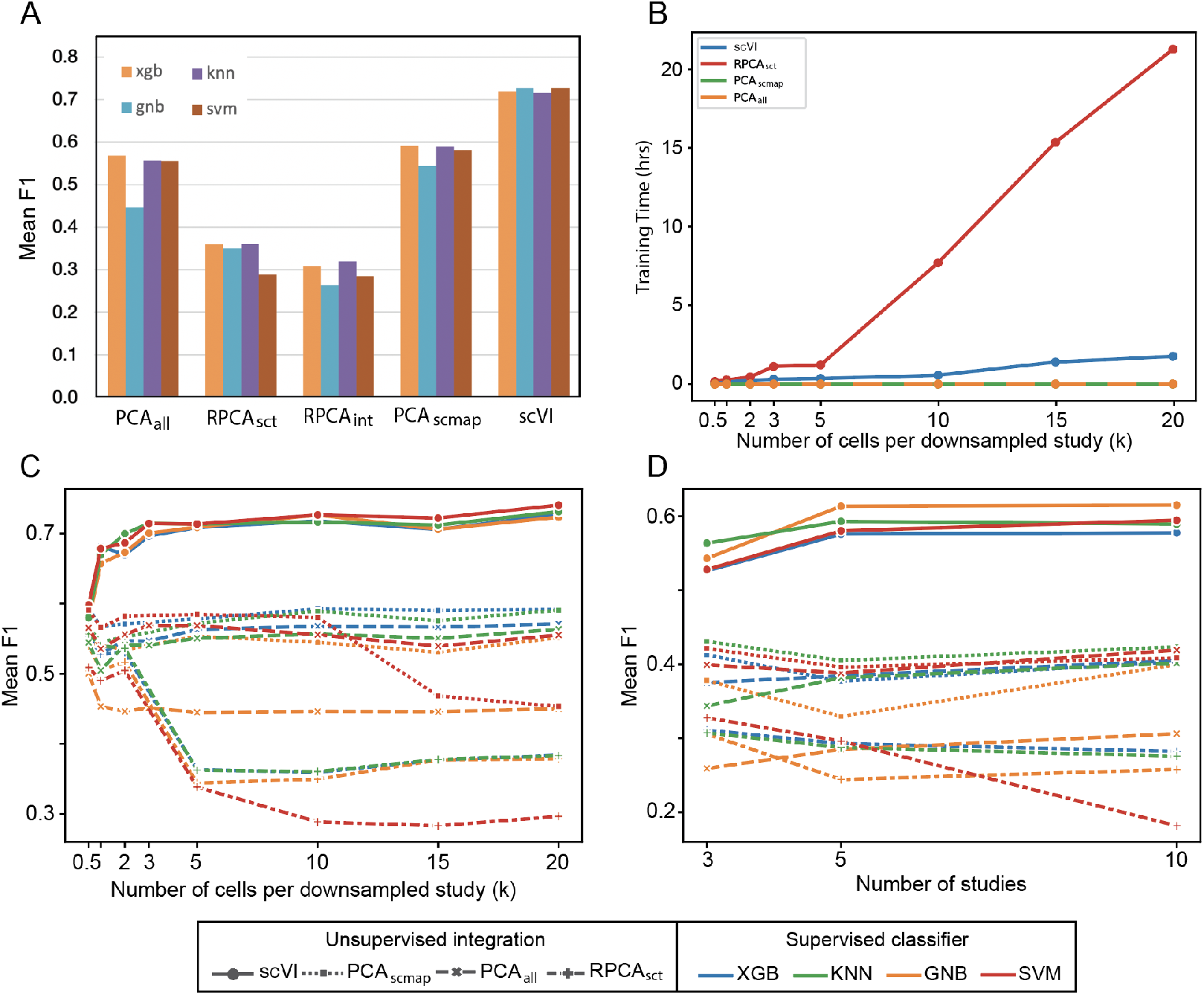
Performance of models on the scMARK benchmark dataset: A) Five feature reduction approaches were compared with four cell-type classifiers. The results highlight that scVI outperformed other approaches based on all supervised models (Table S3 for detailed results); B) Compute times for the unsupervised steps increasing the number of cells per study from 300 to 20,000. The following virtual machines (VM) were used: scVI - GPU VM, 128GB memory; PCA_*all*_ and PCA_*scmap*_ - CPU 4 vCPUs, 160 GiB memory; RPCA_*sct*_ used a CPU VM - 4 vCPUs, 976 GiB memory. C) The mean F1 score as cell-sample size increases is recorded for each unsupervised integration and supervised cell-type classification model pairing. D) Similar to A, but keeping 10,000 cells per study constant while varying the number of training studies (2, 4, or 9). In all tests, the F1 score resulted from leave-one-out cross-validation on the supervised step.

### 3.3 Behavior as dataset size increases

We next assessed how computation time and performance vary with increasing dataset size to understand the scalability of the approaches. We increased samples from 500 cells per dataset (5k total) to 20k cells per dataset (200k total). We measured both compute time and the mean F1 score, as calculated by our collection of supervised models (Fig. 4B & 4C). PCA_*all*_ and PCA_*scmap*_ completed the task in minutes, but the F1 score did not increase. RPCA compute times increased considerably and performance dropped off. In contrast scVI performance improved with increasing dataset size, while train time only increased linearly up to 2hrs for 200k cells. scVI also had reduced memory requirements taking 128 GB memory. The same integration with RPCA_*sct*_ took >600 GB of memory. Overall this analysis demonstrates that scVI was the only tested method to scale and leverage larger training set sizes.

Finally, we tested model performance while increasing the dataset number from 3, to 5, to 10. We measured accuracy on three studies (Azizi et al., 2018, Bassez et al., 2021, and Bi et al., 2021; selected alphabetically) to test the effect of additional training data on performance vs. the addition of datasets that may be easier or more difficult to predict. scVI outperformed the other methods even at lower dataset numbers (Fig. 4D). At five datasets, performance plateaued. Of note, at ten datasets, 300 unique batch IDs exist. We speculate this result leaves room for new models to improve on scVI by making better use of the additional datasets, or better handling large batch numbers.

### 3.4 Characteristics of the unified atlas generated by the top-performing model

Our evaluation framework showed scVI outperformed other methods. To qualitatively assess the results of dataset integrations we used UMAP to project the 10-dimensional embedding into a 2 dimensional space [19].

Coloring the projection by study shows a high level of overlap between all studies (Fig. 5A). Coloring by the standardized author cell-type labels shows a tight grouping between cell-types (Fig. 5B). For example, CD4+ T-cells, CD8+ T-cells, and NK cells form a large leukocyte cluster and have a common lineage, epithelial lineages fall in a similar region of the projection to each other, and Fibroblastic and endothelial cells form independent groups. A critical hypothesis of our evaluation framework is that we can find new cell-types in properly unified cell-type spaces. FOXP3+ T-cells are a well-characterized CD4+ T-cell subtype, implicated in immune suppression, and known as regulatory T-cells (T-regs). Within the T-cell cluster, FOXP3+ expression localizes to a distinct region of the CD4+ T-cell space (Fig. 5C) that contains cells from all ten studies. This result shows that we could find this unlabelled cell subtype by analyzing sub-regions of the CD4+ T-cell cluster. This qualitative assessment, thus, highlights the value of the unified cell-type space generated by scVI.

**Figure 5:**
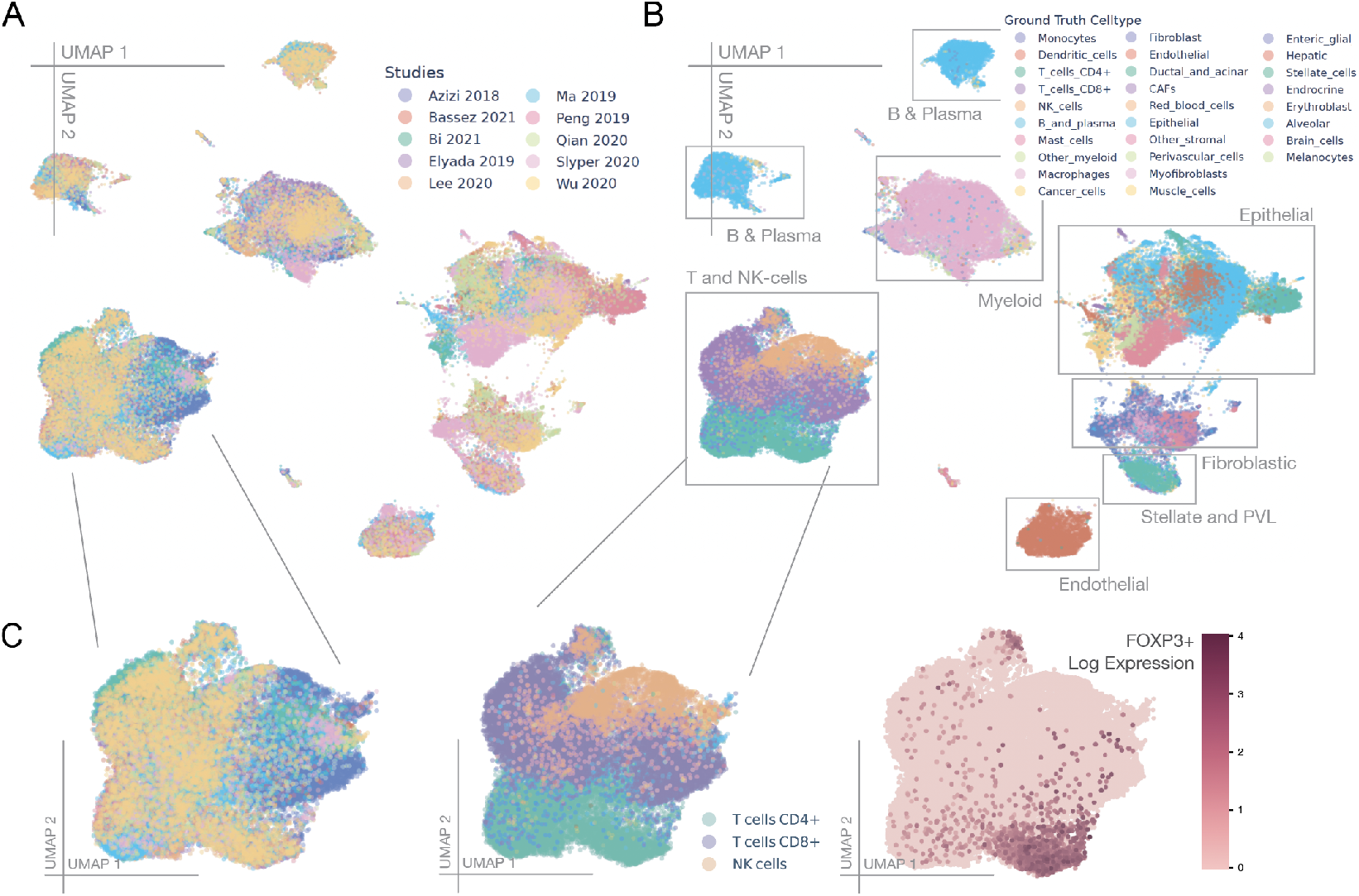
Visual inspection of the results of dataset integration with scVI: A) A UMAP projection of the 10-dimensional embedding space produced by the VAE highlights broad levels of overlap across all studies; B) The same UMAP projection but colored by standardized author cell-type labels (see Table S1 for harmonization details); C) A zoom-in of the T and NK-cell cluster shows FOXP3+ expression localizes to a subregion of CD4+ T-cells; this agrees with existing knowledge about FOXP3+ expression on regulatory T-cells, a CD4+ T-cell subset

### 3.5 Limitations

Finally, we sought to understand which datasets scVI performed well vs. poorly. We generated confusion matrices for three datasets covering the spectrum of scVI integration performance, using the SVM predictions. Firstly, we examined Azizi 2018, a dataset generated on InDrop. All four unsupervised methods perform poorly on this dataset. Examination of the confusion matrix indicates the dataset is limited to immune cells (Fig. 6A). Myeloid sub-types were difficult to separate in this and other datasets (e.g., Qian 2020), indicating this may be an area for improved ground-truth labeling. In contrast, NK, CD4+, and CD8+ T-cells were difficult to separate in Azizi et al. 2018, but easy elsewhere indicating here, scVI is struggling to transfer learnings from the 10X Chromium to the inDrop. In the confusion matrix of Qian et al. 2020, we found models encountered issues splitting cancer cells from epithelial cells and dendritic cells from macrophages (Fig. 6B). These cell-types are of similar cell lineages in both cases, suggesting better models or larger datasets may lead to small improvements. Accuracy was highest on Ma et al. 2019 (Fig. 6C); this is likely due to high-level cell-type labels provided by the authors, showing more detailed cell-type labels create complexity. Overall, models trained on the scVI cell-type space perform well but there is likely room for improvement, both in model quality and developing benchmark datasets with more granular cell-type labels.

**Figure 6:**
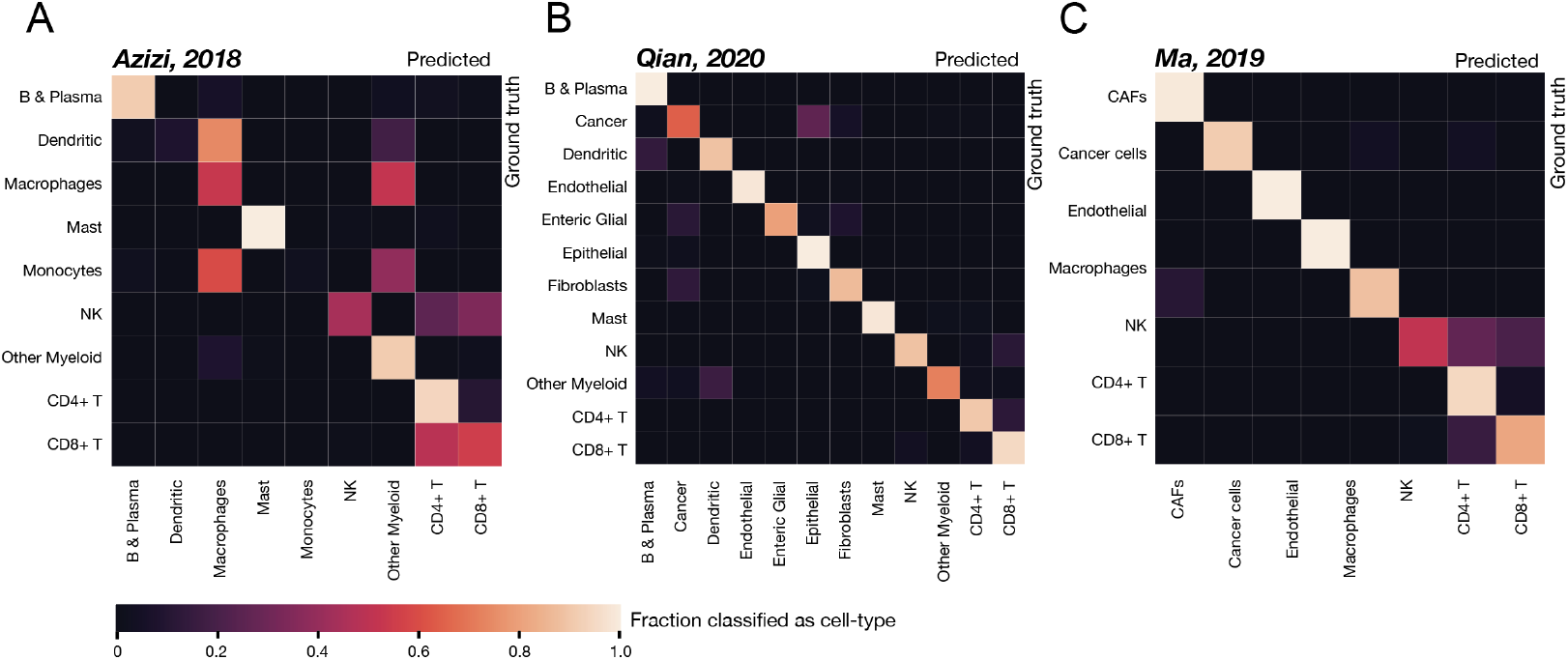
Confusion matrices of scVI embedding with SVM predictions on three scMARK studies: Confusion matrices were generated for held-out datasets using SVM predictions from the scVI cell-type space. Matrices were selected where scVI’s performance was; A) Poor (Azizi 2018; F1 score = 0.483); B) Average (Qian 2020; F1 score = 0.754); and C) Strong (Ma, 2019; F1 score = 0.9).

## 4 Discussion

To date, most benchmarks have used individual studies to test the performance of machine-learning models for analyzing scRNA data. Yet such approaches are a low hurdle. Batch effects within a study are much smaller than those between studies; thus very high-levels of performance are typically seen [4, 5, 20]. Model performance is also commonly tested with statistical metrics [3, 21]. However, how accurately a model aligns cell-types remains an open question under these metrics since there a ground truth is not used. More recent studies use ground-truth labels to test supervised cell-type labeling models on new, unseen studies [4, 22, 23, 7]. Yet, supervised models limit new cell-type discovery; worse, such models can perform well by forcing cell-types into buckets that do not represent their actual state. Collectively these experiments highlight the need for a high-hurdle benchmark that tests models’ ability to align disparate data in a real-world setting.

Here we present, to our knowledge, the first framework and dataset (scMARK) for quantitatively evaluating how well methods can unify scRNA gene-expression matrices into a unified cell-type space that avoids the pitfalls and complexities of using statistical metrics of performance. We achieve this through (1) Creating a benchmark dataset to test how well models can integrate disparate data from different authors; (2) Using an unsupervised scRNA model to integrate the studies into a single unified cell-type space; and (3) Evaluating this space with a collection of supervised models. We show that scVI, a Variational Autoencoder, optimized to manage scRNA batch-effects, outperforms other approaches. We find that scVI also represents the only tested method that benefits from larger training datasets. Qualitative assessment of the unified cell-type space indicates that the scVI embedding is suitable for automatic cell-type labeling and discovery of new cell-types. Thus we find that scVI, and VAE’s more generally represent a solution to scaling unification of scRNA-seq data. We hope this work to develop an MNIST-like scRNA benchmark will pave the way to the generation of larger datasets akin to ImageNet [24]. We speculate robust models derived from larger ImageNet like datasets will, in turn, be critical to both the construction of tissue and disease atlases, and use of scRNA technologies in a clinical setting.

## 5 Methods

### Collection and standardization of scRNA data

Raw scRNA UMI count matrices were obtained from public repositories (Table S1) and quality control followed the original author filters. Gene identifiers were standardized across studies based on (i) Human Protein Atlas (HPA) versions 13 to 20; and (ii) ENSEMBL GRCh38 versions 78 to 103. Priority was given to HPA identifiers. In cases where the authors provided only general T-cell annotations, we used Azimuth [15] to assign those cells into CD8+ or CD4+ T cells.

### scRNA data normalization

For scVI, PCA_*all*_ and PCA_*scmap*_, the count matrices were normalized on a per-cell basis using Scanpy v1.7.2, by dividing each cell by its total count over all genes. The normalized count was then multiplied by a scale factor of 10000, after which a log(X+1) transformation was applied. For RPCA, Seurat’s SCTransform normalization was used with default parameters [15].

### Unsupervised cell-type space methods

For PCA, we experimented with two types of gene selection methods: PCA_*scmap*_ and PCA_*all*_. We used the TruncatedSVD method from sci-kit learn (v0.24.2) library instead of the PCA method because TruncatedSVD is more efficient for sparse matrices. For PCA_*scmap*_, we applied [17] scmap method to the normalized data to extract the top 500 genes, and ran PCA on this. For PCA_*all*_, we applied PCA to the normalized matrix containing all genes.

We reimplemented the scVI variational auto-encoder described by [18]. We used sample level batch-correction and the following hyper-parameters: both encoder and decoder had two linear layers, with 1024 nodes for each linear layer. For both encoder and decoder, dropout regularization (with 0.1 probability of an element to be zeroed) and batch normalization was used in between two hidden layers along with ReLU activation function. The latent space dimension was set to 10 and modeled using the Normal distribution. A zero-inflated negative binomial distribution was used to model gene counts. The Adam optimizer was used for training the VAE with learning rate=5*e* – 4, weight decay=l*e* – 5 and eps=0.01. A step learning rate scheduler was used with a step size=50 epochs and lr factor=0.1, and early stopping with patience=10 epochs. The model was trained with a batch-size of 32 or 64 and for a maximum of 100 epochs. All the implementation was done in Python using Pytorch (1.7.0) library.

The RPCA method was implemented in R using Seurat (v4.0.3) [16], with the top-10 larger samples used as references for anchor detection and the following parameters (dims=10, npcs=10, k.filter=150, k.weight=100). Samples with <150 cells were merged into a single batch with others from the same study and tissue type. For all RPCA analyses we used 4 workers, as a higher number failed to retrieve the result from the MulticoreFuture parallelization function. The output of the functions RunPCA (npcs=10) and RunUMAP (n.components=10), with assay=“SCT” or “integrated” were used as inputs for the supervised cell-type classifiers.

### Supervised cell-type classification methods

Supervised cell-type classification methods: A 90/10 split of the training datasets was made for training/validation and GridSearchCV from the sci-kit learn package (v0.24.2) was used to test and select hyperparameter values (Table S4) based on the best F1-score on the validation set. For XGboost, we used XGBClassifier method with multi-softmax objective function from XGboost (v1.4.0) python library. The remaining cell-type classifiers (GNB, KNN and SVM) were implemented using sci-kit learn. We used GaussianNB, KNeighborsClassifier, and SVC (with output probability) method for GNB, KNN and SVM respectively from sci-kit learn library.

## Acknowledgements

Dr. Chris J. Harvey for helping map cell-type labels. Dr. Sean Grullon for helping initiate the project and in the implementation of scVI.

## Data Availability

scMARK dataset is available at: https://doi.org/10.5281/zenodo.5765804

---

**Table S1.** studies used in benchmark.

| Study ID | Tissues included in scMARK* | Data source** | Reference |
| --- | --- | --- | --- |
| Azizi 2018 | Breast (c & n) | GEO:GSE114725 | Azizi et al (2018) Cell;174(5):1293 |
| Bassez 2021 | Breast (c) | Lambrechts Lab Website | Bassez et al (2021) Nat Med; 27(5):820 |
| Bi 2021 | Kidney (c) | Broad:SCP1288 | Bi et al (2021) Cancer Cell;39(5):649 |
| Elyada 2019 | Pancreas (c & n) | CReSCENT:CRES-P33 | Elyada et al (2019) Cancer Discov;9(8):1102 |
| Lee 2020 | Colorectal (c & n) | GEO:GSE132465 and GEO:GSE144735 | Lee et al (2020) Nat Genet;52(6):594 |
| Ma 2019 | Liver (c) | GEO:GSE125449 | Ma et al (2019) Cancer Cell;36(4):418 |
| Peng 2019 | Pancreas (c & n) | GSA:CRA001160 | Peng et al (2019) Cell Res;29(9):725 |
| Qian 2020 | Breast (c), Colorectal (c & n), Lung (c & n), Ovarian (c & n) | Lambrechts Lab Website | Qian et al (2020) Cell Res;30(9):745 |
| Slyper 2020 | Lung (c), Ovarian (c), Skin (c), Neuroblastoma (c) | GEO:GSE140819 | Slyper et al (2020) Nat Med;26(5):792 |
| Wu 2020 | Breast (c) | Broad:SCP1106 | Wu et al (2020) EMBO J;39(19):e104063 |
| * cancer (c), normal (n) |  |  |  |
| ** Gene Expression Omnibus (GEO), Broad Institute Single Cell Portal (Broad), CanceR Single Cell Expression Toolkit (CReSCENT), Genome Sequence Archive (GSA) |  |  |  |

**Table S2:** Cell type standardization keys.

| B_and_plasma_cells |  |  |  |
| --- | --- | --- | --- |
| B cells | B cell | Plasma cell | B_cell |
| B_Cells | CD19+CD20+ B | IgA+ Plasma | IgG+ Plasma |
| Unspecified Plasma | B: | Plasma cells | Plasma_Cells |

| Dendritic_cells |  |  |  |
| --- | --- | --- | --- |
| LanghDC | CLEC9A+ DC | mDC: | quiescent_migDC |
| pDC | cDC | cDC2 | CD1C+ DC |
| pDC: | Activated DC | DC | cDC1 |
| pDCs | ASDC | mregDC_migDC |  |

| Ductal_and_acinar_cells |  |  |  |
| --- | --- | --- | --- |
| Ductal cells 1 | Ductal cells 2 | Ductal cells 3 | Acinar cell |
| Ductal cell type 2 | Ductal cell type 1 | Acinar cells |  |

| Epithelial |  |  |  |
| --- | --- | --- | --- |
| Myoepithelial | Perivascular cells | Epithelial | BEST4+ Enterocytes |
| Epithelial_Luminal_Mature | Epithelial cell | CMS1 | CMS2 |
| CMS4 | Epithelial cells | Goblet cells | Intermediate |
| Mature Enterocytes | Mature Enterocytes type 1 | Mature Enterocytes type 2 | Tuft cells |
| Stem-like/TA | Epithelial_Basal_Cycling | CMS3 | Epithelial_Basal |

| Macrophages |  |  |  |
| --- | --- | --- | --- |
| C3_CCR2 | Pro-inflammatory | CXCL10-Hi TAM | SPP1+ |
| SPP1+B | VSIR-Hi TAM | C4_CCL2 | C5_CCL18 |
| C6_MMP9 | C9_SLC2A1 | C7_CX3CR1 | C8_MT1G |
| TAM | Macrophage cell | SPP1+A | MACROPHAGE: |
| Cycling TAM | GPNMB-Hi TAM | Anti-inflammatory | FOLR2-Hi TAM |
| Macrophage 2 | Myeloid & Macrophages | Macrophage 1 | Macrophage |
| LowLibSize Macrophage | C10_LYVE1 |  |  |

| T_cells_CD4+ |  |  |  |
| --- | --- | --- | --- |
| CD4+ T-cells | T-cells Cycling | T:CD4+NAIVE | CD4+ T cell |
| T:CD4+CM | T:Reg | T:CD4+EM | CD4_EM |
| CD4_EX | CD4_EX_Proliferating | CD4_N | CD4_REG |
| CD4_REG_Proliferating | Effector T-Helper | Memory T-Helper | MitoHigh T-Helper |
| Tfh cells | CD4+ T cells | T helper 17 cells | T-Reg |
| Regulatory T cells | T-Regs | T follicular helper cells |  |

| T_cells_CD8+ |  |  |  |
| --- | --- | --- | --- |
| MitoHigh CD8+ T cell | T:CD8+EM | T:CD8+NAIVE | T:CD8+CM |
| gamma delta T cells | CD8+ T cells | CD8_EM | CD8+ T-cells |
| CD8_EMRA | CD8_EX_Proliferating | MX1-Hi CD8+ T cell | Cycling CD8+ T cell |
| CD8_EX | 41BB-Hi CD8+ T cell | Vg9Vd2_gdT | 41BB-Lo CD8+ T cell |
| CD8+ T cell | CD8_RM | CD8_N | gdT |

**Table S2:**
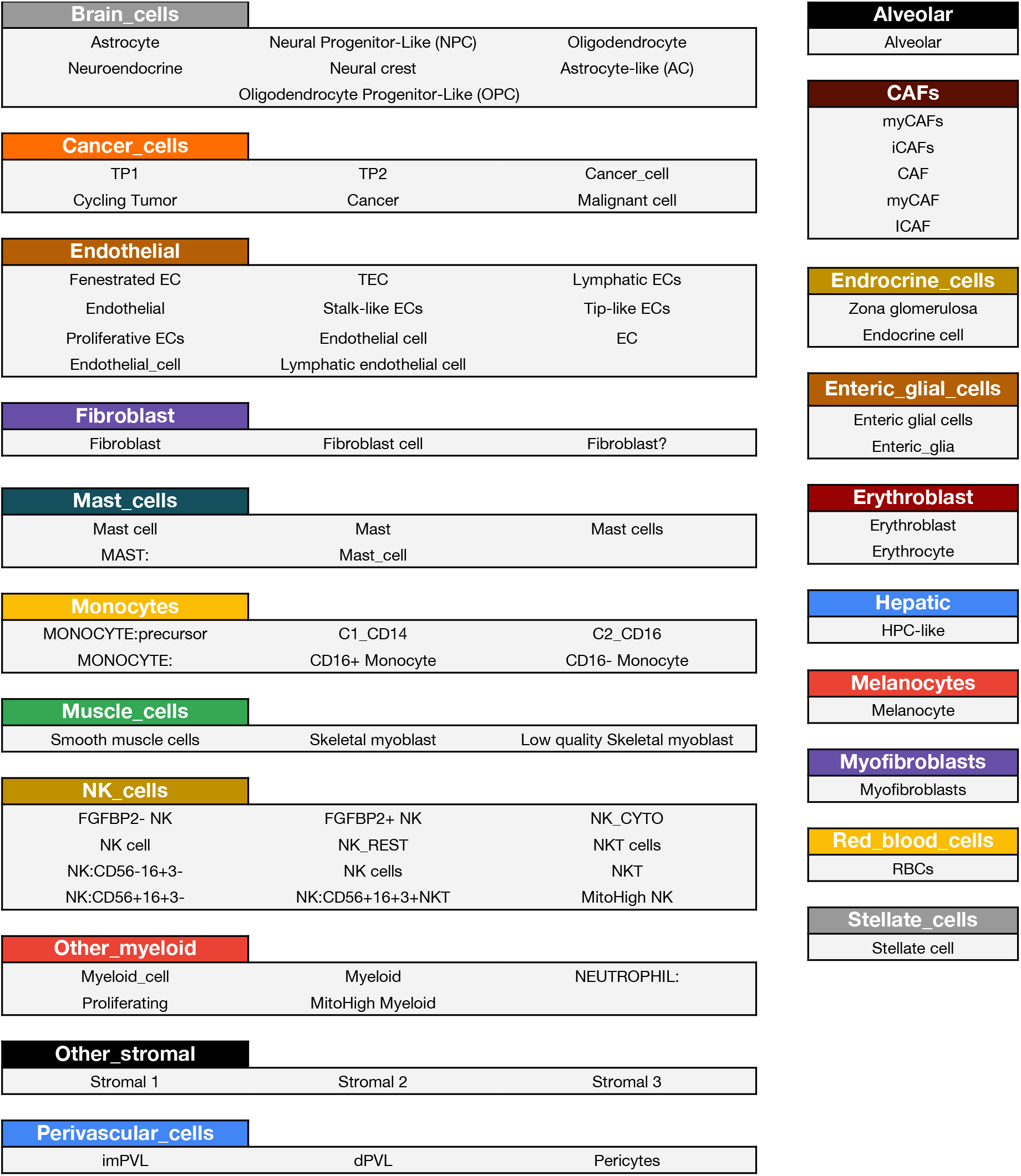
Cell type standardization keys (Cont.)

**Table S3:** Detailed F1 Scores.

| Unsupervised | Supervised | Individual Dataset Performance (F1 Score) |  |  |  |  |  |  |  |  |  | Mean Score |
| --- | --- | --- | --- | --- | --- | --- | --- | --- | --- | --- | --- | --- |
|  |  | Ma, 2019 | Azizi, 2018 | Bi, 2021 | Peng, 2019 | Bassez, 2021 | Elyada, 2019 | Lee, 2020 | Slyper, 2020 | Qian, 2020 | Wu, 2021 |  |
| PCA All | xgb | 0.746 | 0.233 | 0.427 | 0.703 | 0.556 | 0.558 | 0.641 | 0.614 | 0.575 | 0.625 | **0.568** |
|  | gnb | 0.620 | 0.196 | 0.336 | 0.603 | 0.386 | 0.484 | 0.482 | 0.457 | 0.367 | 0.532 | **0.446** |
|  | knn | 0.748 | 0.256 | 0.396 | 0.703 | 0.552 | 0.551 | 0.598 | 0.575 | 0.570 | 0.618 | **0.557** |
|  | svm | 0.758 | 0.288 | 0.397 | 0.700 | 0.572 | 0.580 | 0.497 | 0.545 | 0.579 | 0.640 | **0.556** |
| RPCA | xgb | 0.178 | 0.374 | 0.294 | 0.335 | 0.308 | 0.332 | 0.359 | 0.497 | 0.427 | 0.491 | **0.360** |
|  | nb | 0.210 | 0.296 | 0.267 | 0.298 | 0.309 | 0.360 | 0.346 | 0.535 | 0.367 | 0.508 | **0.350** |
|  | knn | 0.185 | 0.361 | 0.281 | 0.350 | 0.304 | 0.323 | 0.364 | 0.516 | 0.418 | 0.502 | **0.361** |
|  | svm | 0.087 | 0.249 | 0.210 | 0.302 | 0.307 | 0.298 | 0.270 | 0.417 | 0.296 | 0.452 | **0.289** |
| PCA scMap | xgb | 0.221 | 0.507 | 0.482 | 0.619 | 0.575 | 0.605 | 0.663 | 0.798 | 0.713 | 0.743 | **0.593** |
|  | nb | 0.262 | 0.491 | 0.446 | 0.553 | 0.536 | 0.523 | 0.578 | 0.698 | 0.732 | 0.630 | **0.545** |
|  | knn | 0.248 | 0.520 | 0.502 | 0.625 | 0.513 | 0.582 | 0.678 | 0.794 | 0.714 | 0.717 | **0.589** |
|  | svm | 0.249 | 0.529 | 0.447 | 0.638 | 0.582 | 0.622 | 0.552 | 0.788 | 0.719 | 0.676 | **0.580** |
| VAE | xgb | 0.886 | 0.466 | 0.569 | 0.776 | 0.698 | 0.733 | 0.767 | 0.752 | 0.744 | 0.791 | **0.718** |
|  | gnb | 0.887 | 0.511 | 0.615 | 0.745 | 0.719 | 0.737 | 0.800 | 0.746 | 0.703 | 0.802 | **0.727** |
|  | knn | 0.878 | 0.483 | 0.590 | 0.784 | 0.696 | 0.738 | 0.733 | 0.745 | 0.747 | 0.772 | **0.716** |
|  | svm | 0.900 | 0.483 | 0.578 | 0.790 | 0.722 | 0.740 | 0.766 | 0.741 | 0.754 | 0.791 | **0.727** |
| Mean Score |  | 0.504 | 0.390 | 0.427 | 0.595 | 0.521 | 0.548 | 0.568 | 0.639 | 0.589 | 0.643 |  |

**Table S4:** Hyperparameters tested for supervised models.

| Supervised Model | Hyperparameter | Values Searched |
| --- | --- | --- |
| XGB | Eta | {0.1, 0.2, 0.3} |
|  | Gamma | {0.1, 1, 1.5} |
|  | Max depth | {3, 5} |
| SVM | C | 10 |
|  | Kernel | RBF |
| GNB | Smoothing variable | {1e-8, 1e-9, 1e-10} |
| KNN | N neighbors | {5, 9} |

## References

[1] Valentine Svensson, Eduardo da Veiga Beltrame, and Lior Pachter. A curated database reveals trends in single-cell transcriptomics. Database, 2020, 2020.

[2] Zoe A Clarke, Tallulah S Andrews, Jawairia Atif, Delaram Pouyabahar, Brendan T Innes, Sonya A MacParland, and Gary D Bader. Tutorial: guidelines for annotating single-cell transcriptomic maps using automated and manual methods. Nature protocols, 16(6):2749–2764, 2021.

[3] Andrew Butler, Paul Hoffman, Peter Smibert, Efthymia Papalexi, and Rahul Satija. Integrating single-cell transcriptomic data across different conditions, technologies, and species. Nat. Biotechnol., 36(5):411–420, April 2018.

[4] Tamim Abdelaal, Lieke Michielsen, Davy Cats, Dylan Hoogduin, Hailiang Mei, Marcel J T Reinders, and Ahmed Mahfouz. A comparison of automatic cell identification methods for single-cell RNA sequencing data. Genome Biol., 20(1):194, September 2019.

[5] J Javier Diaz-Mejia, J Javier Diaz-Mejia, Elaine C Meng, Alexander R Pico, Sonya A MacParland, Troy Ketela, Trevor J Pugh, Gary D Bader, and John H Morris. Evaluation of methods to assign cell type labels to cell clusters from single-cell RNA-sequencing data, 2019.

[6] Qianhui Huang, Yu Liu, Yuheng Du, and Lana X Garmire. Evaluation of cell type annotation R packages on single-cell RNA-seq data. Genomics Proteomics Bioinformatics, 19(2):267–281, April 2021.

[7] Bingbing Xie, Qin Jiang, Antonio Mora, and Xuri Li. Automatic cell type identification methods for single-cell RNA sequencing. Comput. Struct. Biotechnol. J., 19:5874–5887, October 2021.

[8] Malte D Luecken, Daniel Bernard Burkhardt, Robrecht Cannoodt, Christopher Lance, Aditi Agrawal, Hananeh Aliee, Ann T Chen, Louise Deconinck, Angela M Detweiler, Alejandro A Granados, et al. A sandbox for prediction and integration of dna, rna, and proteins in single cells. In Thirty-fifth Conference on Neural Information Processing Systems Datasets and Benchmarks Track (Round 2), 2021.

[9] Y Lecun, L Bottou, Y Bengio, and P Haffner. Gradient-based learning applied to document recognition. Proc. IEEE, 86(11):2278–2324, November 1998.

[10] Tianyu Wang, Boyang Li, Craig E Nelson, and Sheida Nabavi. Comparative analysis of differential gene expression analysis tools for single-cell RNA sequencing data. BMC Bioinformatics, 20(1):40, January 2019.

[11] Hoa Thi Nhu Tran, Kok Siong Ang, Marion Chevrier, Xiaomeng Zhang, Nicole Yee Shin Lee, Michelle Goh, and Jinmiao Chen. A benchmark of batch-effect correction methods for single-cell RNA sequencing data. Genome Biol., 21(1):12, January 2020.

[12] Z Ren, M Gerlach, H Shi, G R S Budinger, and L A N Amaral. Information-theory-based benchmarking and feature selection algorithm improve cell type annotation and reproducibility of single cell RNA-seq data analysis… bioRxiv, 2020.

[13] Xiannian Zhang, Tianqi Li, Feng Liu, Yaqi Chen, Jiacheng Yao, Zeyao Li, Yanyi Huang, and Jianbin Wang. Comparative analysis of Droplet-Based Ultra-High-Throughput Single-Cell RNA-Seq systems. Mol. Cell, 73(1):130–142.e5, January 2019.

[14] Mathias Uhlen, Max J Karlsson, Wen Zhong, Abdellah Tebani, Christian Pou, Jaromir Mikes, Tadepally Lakshmikanth, Björn Forsström, Fredrik Edfors, Jacob Odeberg, et al. A genome-wide transcriptomic analysis of protein-coding genes in human blood cells. Science, 366(6472), 2019.

[15] Kevin L Howe, Premanand Achuthan, James Allen, Jamie Allen, Jorge Alvarez-Jarreta, M Ridwan Amode, Irina M Armean, Andrey G Azov, Ruth Bennett, Jyothish Bhai, et al. Ensembl 2021. Nucleic acids research, 49(D1):D884–D891, 2021.

[16] Yuhan Hao, Stephanie Hao, Erica Andersen-Nissen, William M Mauck, 3rd, Shiwei Zheng, Andrew Butler, Maddie J Lee, Aaron J Wilk, Charlotte Darby, Michael Zager, Paul Hoffman, Marlon Stoeckius, Efthymia Papalexi, Eleni P Mimitou, Jaison Jain, Avi Srivastava, Tim Stuart, Lamar M Fleming, Bertrand Yeung, Angela J Rogers, Juliana M McElrath, Catherine A Blish, Raphael Gottardo, Peter Smibert, and Rahul Satija. Integrated analysis of multimodal single-cell data. Cell, 184(13):3573–3587.e29, June 2021.

[17] Vladimir Yu Kiselev, Andrew Yiu, and Martin Hemberg. scmap: projection of single-cell RNA-seq data across data sets. Nat. Methods, 15(5):359–362, May 2018.

[18] Romain Lopez, Jeffrey Regier, Michael B Cole, Michael I Jordan, and Nir Yosef. Deep generative modeling for single-cell transcriptomics. Nat. Methods, 15(12):1053–1058, December 2018.

[19] Leland McInnes, John Healy, and James Melville. UMAP: Uniform manifold approximation and projection for dimension reduction. February 2018.

[20] Giovanni Pasquini, Jesus Eduardo Rojo Arias, Patrick Schäfer, and Volker Busskamp. Automated methods for cell type annotation on scRNA-seq data. Comput. Struct. Biotechnol. J., 19:961–969, January 2021.

[21] M D Luecken, M Büttner, K Chaichoompu, A Danese, and others. Benchmarking atlas-level data integration in single-cell genomics. BioRxiv, 2020.

[22] Xinlei Zhao, Shuang Wu, Nan Fang, Xiao Sun, and Jue Fan. Evaluation of single-cell classifiers for single-cell RNA sequencing data sets. Brief. Bioinform., 21(5):1581–1595, September 2020.

[23] Yixuan Huang and Peng Zhang. Corrigendum to: Evaluation of machine learning approaches for cell-type identification from single-cell transcriptomics data. Brief. Bioinform., 22(6), November 2021.

[24] Jia Deng, Wei Dong, Richard Socher, Li-Jia Li, Kai Li, and Li Fei-Fei. ImageNet: A large-scale hierarchical image database. In 2009 IEEE Conference on Computer Vision and Pattern Recognition, pages 248–255, June 2009.

[25] Elham Azizi, Ambrose J Carr, George Plitas, Andrew E Cornish, Catherine Konopacki, Sandhya Prabhakaran, Juozas Nainys, Kenmin Wu, Vaidotas Kiseliovas, Manu Setty, et al. Single-cell map of diverse immune phenotypes in the breast tumor microenvironment. Cell, 174(5):1293–1308, 2018.

[26] Ayse Bassez, Hanne Vos, Laurien Van Dyck, Giuseppe Floris, Ingrid Arijs, Christine Desmedt, Bram Boeckx, Marlies Vanden Bempt, Ines Nevelsteen, Kathleen Lambein, et al. A single-cell map of intratumoral changes during anti-pd1 treatment of patients with breast cancer. Nature Medicine, 27(5):820–832, 2021.

[27] Ela Elyada, Mohan Bolisetty, Pasquale Laise, William F Flynn, Elise T Courtois, Richard A Burkhart, Jonathan A Teinor, Pascal Belleau, Giulia Biffi, Matthew S Lucito, et al. Cross-species single-cell analysis of pancreatic ductal adenocarcinoma reveals antigen-presenting cancer-associated fibroblasts. Cancer discovery, 9(8):1102–1123, 2019.

[28] Hae-Ock Lee, Yourae Hong, Hakki Emre Etlioglu, Yong Beom Cho, Valentina Pomella, Ben Van den Bosch, Jasper Vanhecke, Sara Verbandt, Hyekyung Hong, Jae-Woong Min, et al. Lineage-dependent gene expression programs influence the immune landscape of colorectal cancer. Nature Genetics, 52(6):594–603, 2020.

[29] Lichun Ma, Maria O Hernandez, Yongmei Zhao, Monika Mehta, Bao Tran, Michael Kelly, Zachary Rae, Jonathan M Hernandez, Jeremy L Davis, Sean P Martin, et al. Tumor cell biodiversity drives microenvironmental reprogramming in liver cancer. Cancer cell, 36(4):418–430, 2019.

[30] Junya Peng, Bao-Fa Sun, Chuan-Yuan Chen, Jia-Yi Zhou, Yu-Sheng Chen, Hao Chen, Lulu Liu, Dan Huang, Jialin Jiang, Guan-Shen Cui, et al. Single-cell rna-seq highlights intra-tumoral heterogeneity and malignant progression in pancreatic ductal adenocarcinoma. Cell research, 29(9):725–738, 2019.

[31] Junbin Qian, Siel Olbrecht, Bram Boeckx, Hanne Vos, Damya Laoui, Emre Etlioglu, Els Wauters, Valentina Pomella, Sara Verbandt, Pieter Busschaert, et al. A pan-cancer blueprint of the heterogeneous tumor microenvironment revealed by single-cell profiling. Cell research, 30(9):745–762, 2020.

[32] Michal Slyper, Caroline BM Porter, Orr Ashenberg, Julia Waldman, Eugene Drokhlyansky, Isaac Wakiro, Christopher Smillie, Gabriela Smith-Rosario, Jingyi Wu, Danielle Dionne, et al. A single-cell and single-nucleus rna-seq toolbox for fresh and frozen human tumors. Nature medicine, 26(5):792–802, 2020.

[33] Sunny Z Wu, Daniel L Roden, Chenfei Wang, Holly Holliday, Kate Harvey, Aurélie S Cazet, Kendelle J Murphy, Brooke Pereira, Ghamdan Al-Eryani, Nenad Bartonicek, et al. Stromal cell diversity associated with immune evasion in human triple-negative breast cancer. The EMBO Journal, 39(19):e104063, 2020.

